# Time-resolved decoding of uncertainty about stimulus features from human EEG

**DOI:** 10.64898/2026.09.08.750188

**Authors:** Jeffrey Nestor, Karen J. Tian, Angus F. Chapman, Rachel N. Denison

## Abstract

Internal and external noise produce uncertainty in the neural representations of sensory stimuli. Uncertainty about basic stimulus features can be decoded from responses in sensory brain regions by estimating full probability distributions over stimulus features, rather than point estimates. Such decoded uncertainty correlates with subjective uncertainty reports, providing insight into the neural basis of metacognitive judgments. However, in humans, probabilistic decoding has only been applied to functional magnetic resonance imaging (fMRI) data, which has low temporal resolution and thus can give only limited insight into the dynamics of uncertainty in the brain. Here, we assessed whether probabilistic decoding of uncertainty about stimulus features could be extended to electroencephalography (EEG) data, which has higher temporal resolution but lower spatial resolution and different noise properties compared to fMRI. Participants performed a spatial location estimation task and provided subjective uncertainty reports. We found that time-resolved probabilistic decoding in EEG was feasible, as decoders produced accurate predictions of stimulus location following stimulus onset, and decoding error correlated trial-by-trial with decoded uncertainty. The choice of noise covariance structure critically impacted these metrics. Further, decoded uncertainty was a more reliable trial-by-trial indicator of stimulus information than decoding error, illustrating the advantages of probabilistic decoding over standard decoding approaches. However, uncertainty decoded from EEG had no significant trial-by-trial correlation with subjective uncertainty at any time point. Based on these results, uncertainty decoded from EEG provides a time-resolved estimate of stimulus information available from the brain signal on a single trial but may not relate to metacognitive reports.

## Introduction

Human observers are constantly faced with unreliability in the sensory information available to them. For instance, when driving in the fog, you may be uncertain of the distance to the next car, which should lead you to drive more cautiously. In addition to external sources of uncertainty such as fog, it is advantageous if an observer can also account for uncertainty generated internally by neural variability (Tolhurst et al., 1983). Indeed, humans have insight into their level of uncertainty regardless of its source: the subjective uncertainty people report covaries with their objective performance, even across repeated presentations of the same stimulus (Barthelmé & Mamassian, 2009). This metacognitive sensitivity facilitates various aspects of behavior, including efficient communication (Bahrami et al., 2010) and strategic information seeking (Desender et al., 2018), and it is disrupted in mental health conditions including schizophrenia (David et al., 2012) and major depressive disorder (Hong et al., 2025).

In addition to its ramifications for metacognition and decision-making, uncertainty may play a pervasive role in unconscious perceptual processing. According to assertions that perception is a process of Bayesian inference (Born & Bencomo, 2021), the brain must track uncertainty at each stage of processing to determine the degree to which sensory evidence should impact perception. The neural mechanisms underlying such unconscious uncertainty representations, and whether or not the same mechanisms are also involved in the explicit readout of subjective uncertainty, remain unclear.

A promising approach to investigating the neural representations of uncertainty involves directly quantifying the uncertainty about external variables inherent to the measured brain signal based on an inferred neural code. For example, given a set of brain responses, known properties of neural tuning can be used to decode a probability distribution over some stimulus feature, and the variance of that distribution can be used as a measure of uncertainty (van Bergen et al., 2015; Walker et al., 2023). Such code-driven work has shown that decoded uncertainty can be related to human behavior, including subjective uncertainty judgments (Geurts et al., 2022; Li et al., 2021). However, in humans, this approach has only been applied to fMRI data, which has low temporal resolution, and so cannot reveal the dynamics of uncertainty over subsecond timescales relevant for perception and cognition.

Studies of uncertainty representations using high-temporal-resolution neuroimaging modalities have identified neural responses that correlate with behavioral measures like reported confidence (Desender et al., 2019; Dou et al., 2024; Stone et al., 2024). However, such a correlational approach cannot capture the uncertainty associated with sensory encoding (Walker et al., 2023). A code-driven approach to studying uncertainty has not yet been developed for human neuroimaging methods with high temporal resolution such as EEG. It is unclear whether the uncertainty decoding methods developed for fMRI would successfully translate to EEG because of the different spatial resolutions and noise properties of these modalities (van Bergen et al., 2015).

The ability to measure the dynamics of decoded uncertainty in humans could open new research avenues. For example, it could allow investigation of how the brain generates uncertainty about dynamic stimuli, how it incorporates prior information across time, and how feedforward vs. feedback signals contribute to behavioral uncertainty (Ma et al., 2006; Zhu et al., 2024). Uncertainty decoding also has the potential to provide single-trial estimates of time-varying stimulus information, unlike traditional decoding methods that produce only a single decoding accuracy timeseries across all trials.

Here, we developed a method for time-resolved decoding of uncertainty by adapting an existing uncertainty decoding algorithm known as TAFKAP (van Bergen & Jehee, 2021) for EEG. We recorded EEG while human observers completed a visual location estimation task in which targets appeared at iso-eccentric locations around fixation and participants reported the polar location of the target as well as their location uncertainty via a wager task. Then we applied a probabilistic decoding algorithm that computed probability distributions over possible stimulus locations at each time point, providing a time-resolved measure of decoded uncertainty for each trial. To evaluate the algorithm’s performance, we compared decoded uncertainty to decoding error, behavioral error, and reported uncertainty. We also tested variants of the algorithm to accommodate the distinct noise structure of EEG data. We have integrated these analysis methods into an open-source Probabilistic Decoding Across Time (PDAT) toolbox to facilitate future research into the dynamics of uncertainty.

## Methods

### Participants

Fifteen participants (9 female, mean age = 26.4 years) including authors JAN, KJT and AFC were included in the final dataset. This sample size is comparable to other studies using both EEG decoding (Bai et al., 2026; Gurariy et al., 2022) and probabilistic decoding (Li et al., 2021; van Bergen & Jehee, 2019). Data from two other participants were not included because they had fewer than 250 trials of usable data when the target sample size of 15 was reached. All participants had normal or corrected-to-normal vision and were monetarily compensated for their time. All participants provided informed consent and Boston University’s Institutional Review Board approved the experimental protocols.

### Stimuli

Stimuli were displayed on a VIEWPixx/EEG 120 Hz Display (VPixx Technologies Inc., Saint-Bruno, Canada) at a viewing distance of 68 cm on a 50% gray background (37 cd/m^2^). Targets were sinusoidal radial gratings (2 degrees of visual angle (dva) diameter, 4 cycles/° spatial frequency, with a circular aperture aligned with a 50% gray point in the sinusoidal rings). Targets appeared with uniform probability at iso-eccentric locations (72 locations evenly spaced every 5 degrees, 8 dva eccentricity) around fixation. Target contrast was thresholded prior to data collection so that each participant’s mean absolute error in orientation estimation was approximately 10° (thresholded contrast mean: 12%, standard deviation: 3%). Throughout each trial, a central fixation dot (0.3 dva diameter, white with a 0.02 dva black border) was centered on the screen. Additionally, two 75% grey rings (0.05 dva line width) were centered on fixation, with radii of 6 and 10 dva, so that targets would appear between the two rings with a 1 dva buffer on either side (Figure 1). Precues were white rings (0.05 dva line width), centered on fixation with diameter 1.25 dva. The response dot (0.5 dva diameter) was blue ([0, 0, 255]) until participants submitted their location response, at which point it turned black. The response arc (0.2 dva line width) was blue ([0, 0, 255]) until participants submitted their uncertainty response, at which point it turned white. The response dot and arc remained visible during the presentation of the true location feedback, which consisted of a green ([0, 255, 0]) dot (0.5 dva diameter) centered at the same location as the target.

**Figure 1.**
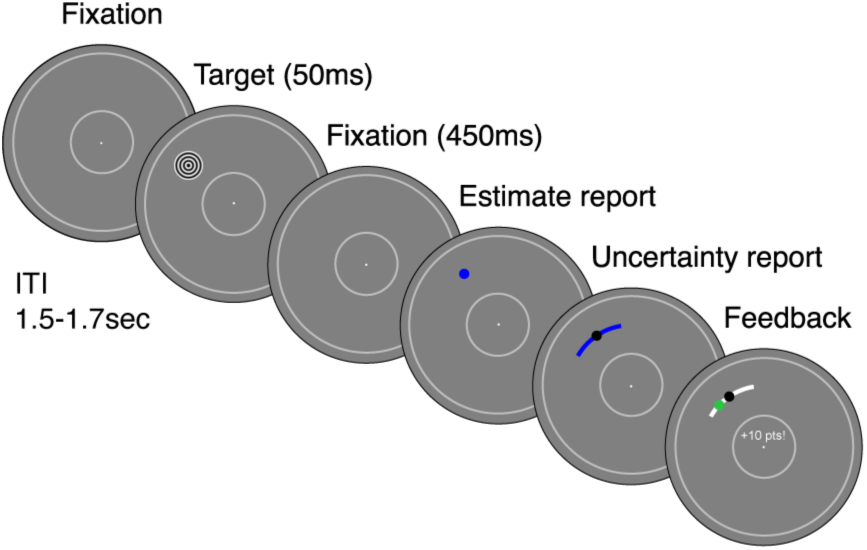
Task schematic for spatial location estimation with uncertainty report. Targets were iso–eccentric radial gratings, with contrast thresholded for each participant to yield approximately 10^∘^ average absolute error. Participants reported location uncertainty by adjusting the width of an arc. More points were awarded for smaller arc lengths, but points were only awarded if the center of the target fell within the arc.

**Figure 2.**
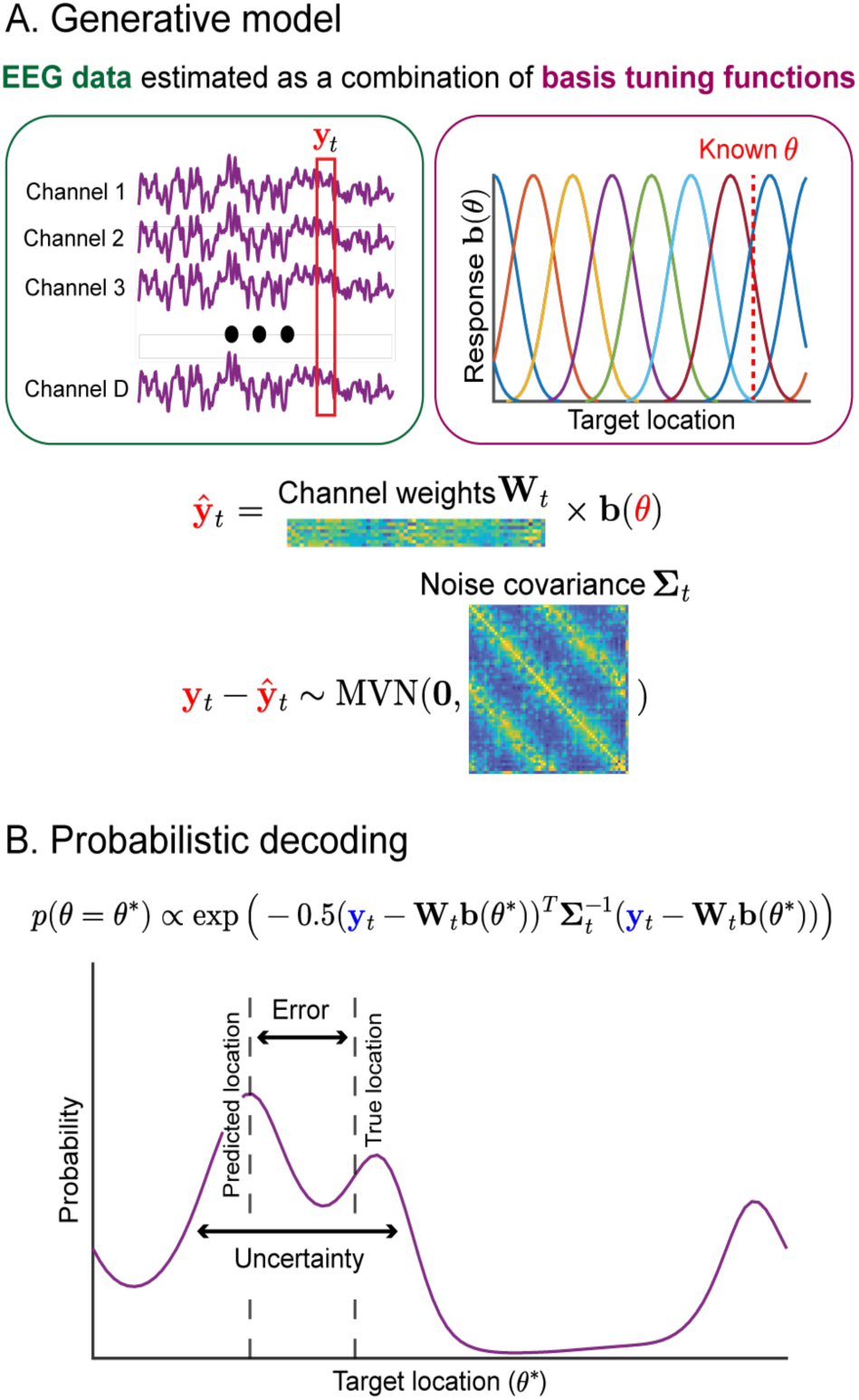
Schematic pipeline of decoding method. A) Generative model. The model assumes that EEG data is the sum of a spatial tuning signal given by a linear combination of *K* basis functions and a noise term. Red symbols refer to an arbitrary trial in the training dataset. Training data is used to fit two key parameters: the weight matrix **W***_t_* (*D×K*), which describes the contribution of each basis function to each EEG channel, and the noise covariance **Σ***_t_* (*D×D*), which describes the distribution of the noise term. B) Probabilistic decoding. Blue symbols refer to an arbitrary test trial. To obtain a posterior probability distribution over target location conditioned on the data from a given test trial, relative likelihoods are computed for 100 evenly spaced locations and then normalized.

### Task

Participants completed a spatial location estimation task on visual targets appearing on a circle around fixation. Each participant completed 432 trials of the main task condition over the course of two sessions on different days, with breaks every 48 trials. Participants for whom fewer than 250 trials remained after excluding trials with fixation breaks, blinks, and poor EEG data quality were invited to complete a third session, bringing their total trial count to 648, until the target participant sample size was reached. In the final dataset, 11 participants completed two sessions and 4 participants completed three sessions, with 294-533 included trials per participant.

Each trial was preceded by a variable inter-trial interval between 1.5 and 1.7 seconds, uniformly distributed in 50 ms increments. The start of the trial was signaled by a central precue (50 ms duration) that provided no information about the upcoming target location. A separate condition was interleaved that contained predictive central precues, but these additional trials were not considered in the analyses reported here. The target stimulus (50 ms duration) followed the precue with a variable stimulus onset asynchrony (SOA) uniformly distributed between 250 and 450 ms. After a 500 ms delay following target onset, a response dot appeared in the center of the screen, signaling that participants could report their location estimate. Once the participant moved the mouse, the response dot jumped from the center of the screen to the target eccentricity, in the direction of the mouse movement. Participants moved the mouse left and right to move the response dot around the ring and clicked to submit their location estimate. To report location uncertainty, participants then used the mouse to adjust the width of an arc around their estimate (Honig et al., 2020; Li et al., 2021) (Figure 1). They received points only if the true location of the center of the target was within their arc, and they received more points the smaller their arc was, incentivizing them to wager on their sense of location uncertainty. The possible point value for an arc that spanned *x*° was equal to 100*e*^(0.008*x*)^ rounded to the nearest integer (Li et al., 2021). After clicking to submit their arc, participants were shown the true location of the target and informed of points received. Additionally, during this feedback stage, participants were informed if any fixation breaks or blinks were detected during the trial. Fixation breaks and blinks were defined as deviations of at least 2.5 dva from fixation and lapses in pupil tracking, respectively.

### Training and thresholding

In a separate session prior to the main task, participants received training on the task and underwent a thresholding procedure to determine the target contrast. The training protocol had three stages. First, participants were familiarized with the basic task structure by completing trials with maximum contrast targets and no eye monitoring. In the second stage, eye monitoring was introduced. In the third stage, target contrast was reduced to 0.15. In each stage, participants completed trials until they felt comfortable with the task. After completing the training, participants underwent a contrast thresholding procedure using the method of constant stimuli. Participants viewed target gratings and submitted location estimates with conditions matched to the main task, except without uncertainty reports or feedback. Location estimates, on targets which tiled the circle, were collected for each of six contrast levels (log contrast: −1.5, −1.3, −1.1, −0.9, −0.7, −0.5), with 24 trials per contrast, for a total of 144 trials. Then, a logistic psychometric function was fit to the mean absolute error per contrast level using MATLAB’s *fminsearch* function. We used this psychometric function to estimate the contrast that would correspond to 10° of absolute error, and this contrast was used for all trials in the main task for that participant.

### Data collection

Participants’ heads were stabilized using a chin-and-head rest. Throughout the experiment, we recorded EEG data using a 64-channel BrainVision actiCHamp system (Brain Vision LLC, Morrisville, NC). The scalp electrodes were referenced online to an electrode on the right mastoid. The recordings were sampled at 1000 Hz with no online filtering. Electrode gel was used to keep impedances below 25 kΩ. To avoid artifacts from blinks and eye movements, participants were instructed to keep their eyes on the fixation stimulus and try not to blink during the period between the appearance of the precue and the response dot.

Gaze position was recorded with an Eyelink SR1000 eye-tracker (SR Research, Ottawa, Canada). Raw gaze positions were converted into degrees of visual angle using five-point grid calibration, which was performed at the start of each session.

### Behavioral analysis

To determine whether participants exhibited metacognitive sensitivity in this task, we examined the trial-by-trial correlation between arc length (from the uncertainty report) and response error (from the location report) for each participant. In addition to assessing this correlation in the original behavioral dataset, we created an adjusted dataset in which we accounted for two factors that could have impacted correlation values. First, to ensure correlations were not driven entirely by “miss trials,” in which participants did not see the stimulus at all and as a result had high response error and high uncertainty, we fit a mixture model where the participant’s location estimate *θ*_response_ has the distribution:

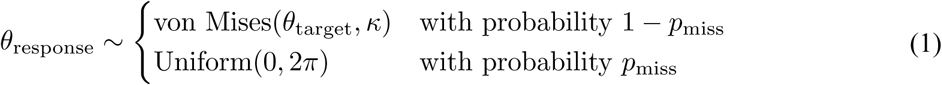

For each participant, the parameters *κ* and *p*_miss_ were fit using MCMC methods, and then the *p*_miss_ portion of trials with the highest absolute response error were excluded from the adjusted dataset. This procedure excludes trials with a high likelihood of being misses, though it may misclassify a small proportion of trials. Then, to ensure correlations were not driven by time-on-task effects, which might occur if, for example, participants’ error and uncertainty both decreased over the course of the experiment as they gained practice with the task, we fit linear regressions to error and uncertainty as a function of trial number, and performed the rest of the analyses on the residual error and uncertainty after subtracting out the fitted time-on-task effect.

#### Statistics

For both the original and corrected datasets, we took the Pearson correlation between arc length and absolute response error for each participant. We then Fisher z-transformed the correlation coefficients and used a z-test to compare them to zero (2-tailed, α = 0.05).

### EEG preprocessing

EEG data were rereferenced offline to the average of the left and right mastoids, and epochs spanning - 200 ms to 500 ms after stimulus onset were extracted from each trial. Any consistently noisy or flat channels were identified manually and interpolated (1 channel each for 3 participants). We used eyetracking data to identify trials with fixation breaks or blinks within this time window and excluded them from analysis. Fixation breaks were defined as deviations greater than 2.5 dva from the center of the screen, and blinks were defined as times when recorded pupil size fell below the 1.5% quantile of pupil size for that session. We also used ERPLAB’s peak-to-peak threshold function to identify and exclude trials with other artifacts (mean proportion of trials excluded due to either eye movements or EEG artifacts = 18%). Epochs were baseline-corrected by subtracting each electrode’s mean amplitude between −200 ms and 0 ms. For the decoding analysis, no offline filtering was performed. Instead, we computed 10 ms sliding window averages and pattern z-scored electrode amplitudes so that the mean value across electrodes on each timepoint in each trial was zero. Because excluded trials had various target locations, we subsampled trials for each participant so that each quadrant of the stimulus space was equally represented in the training set to avoid biasing the model to particular spatial locations.

### EEG decoding

To decode the stimulus location and location uncertainty, we adapted TAFKAP (van Bergen & Jehee, 2021), a probabilistic decoding method based on a generative model in which activity at each recording channel is a noisy function of stimulus location. The location tuning function and the multivariate noise distribution were fit to each participant’s training dataset, and then the generative model was inverted for held-out test trials to compute the posterior probability that the target was in any given location.

#### Generative model

Suppose that, for a given participant, the EEG response at time point *t*, **y***_t_* (*D×*1), to a stimulus at angular location *θ* is given by:

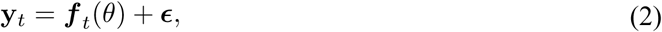

where ***f****_t_* (*θ*) (*D×*1) is a vector containing the fitted tuning functions for each EEG channel at that time point. Each EEG tuning function is based on a linear combination of *K* bell-shaped basis tuning functions:

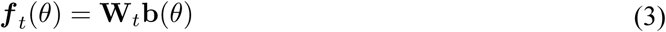

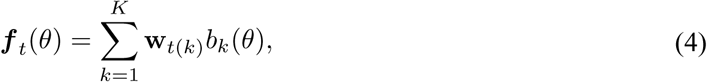

where the weights **W***_t_* (*D×K*) represent the time-varying contribution of each tuning function to each EEG channel’s response and where *b_k_* (*θ*) is the output of the *k*-th tuning function. Each tuning function is defined as a half-wave rectified cosine function raised to the fifth power:

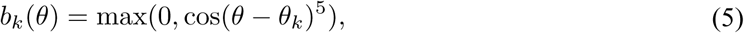

where the *θ_k_*’s evenly tile the circular space [0, 2π].

The noise term **ɛ** follows a multivariate normal distribution whose covariance structure **Σ (***D×D*), describing correlated noise across EEG channels, is allowed to change based on time within the trial:

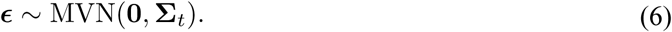

The posterior probability density at a particular stimulus value *θ*\*, conditioned on the EEG response observed at a given time point on a given trial, can be expressed according to Bayes’ rule as:

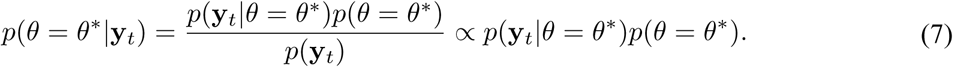

Here we assume a uniform prior across locations: 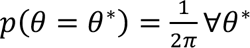. To derive the likelihood *p*(**y***_t_*|*θ* = *θ*^∗^), observe that (2) and (6) imply:

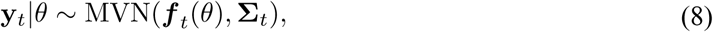

so we can write the probability density function:

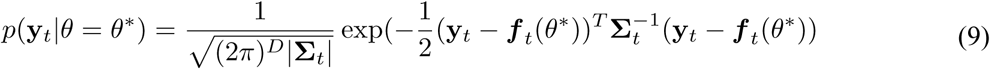

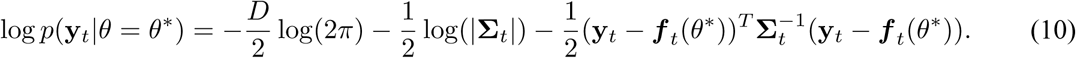

Here we assume that **Σ** does not depend on *θ*, so:

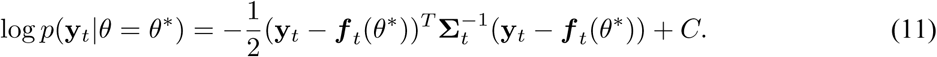

To attain a posterior probability distribution *p*(*θ*|**y***_t_*), we compute the above log likelihood for 100 evenly spaced values of *θ*\*, then exponentiate, and finally normalize the results. Since exponentiation turns the constant term *C* into a multiplier, it cancels out during normalization, so it can be ignored.

#### Fitting the model

The model is fit independently for each participant and each time point, and consists of (1) an estimate of the participant’s *D* tuning functions (one per EEG channel) at that time point, encoded in the *D×K* weight matrix **W***_t_*, as well as (2) an estimate of the *D×D* noise covariance matrix at that time point. In order to reduce model variance, for each iteration of the cross-validation loop, we used a bagging procedure where training trials were resampled with replacement to produce 100 different training sets. Additionally, within each bagging iteration, we randomly selected one of 4 sets of basis functions, each identical up to rotation, in order to avoid model bias due to heterogeneity in the representation of each location by the set of possible tuning functions.

##### 1 Estimating the channel tuning functions

The hypothetical basis tuning functions are set beforehand by the experimenter, so we only need to estimate the weights **w** of each basis tuning function on each channel response. This estimation is done using an ordinary least squares approach. The responses **b**(*θ*) of the *K* basis tuning functions are computed for all *N* training trials given the presented stimulus locations (see Eq. 5) and are encoded in an *N×K* matrix **B**. For each time point, the matrix **Y***_t_* (*N×D*) gives the measured channel responses for all training trials, and channel weights are encoded in the *D×K* matrix **W***_t_*.

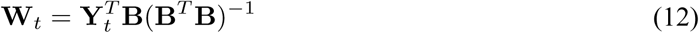

Here, we arbitrarily set the number of basis tuning functions *K* to 8, as we ran preliminary tests with larger *K* and found that it did not impact decoding.

##### 2 Estimating the noise covariance

Sample noise covariance is an unbiased estimate of the true noise covariance:

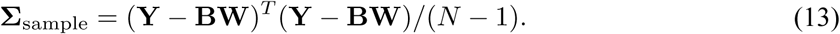

However, **Σ**_sample_ can be a high-variance estimator, including in the EEG noise regime (Blankertz et al., 2011), tending to exhibit eigenvalues more extreme than the true underlying covariance. To reduce variance, it can be advantageous to use an alternative covariance estimator. For most of our analyses, we used the standard shrinkage estimator, which is a common method for EEG data and does not require computationally expensive cross-validation. However, we also tested two other covariance estimators for a key time window to investigate whether more specialized covariance structures may contribute performance benefits in the context of probabilistic decoding of EEG. Finally, we tested an independent variance model and an identity matrix model to characterize the overall importance of the noise covariance in probabilistic decoding of EEG.

##### 2a Standard shrinkage

In the standard shrinkage approach, eigenvalues are regularized by bringing off-diagonal elements closer to zero and elements on the diagonal closer to the mean eigenvalue *ν* of **Σ**_sample_:

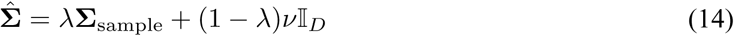

where I*_D_* is the *D×D* identity matrix. Here, we chose *λ* which minimizes the expected squared Froebenius distance between the shrunken covariance matrix and the actual covariance:

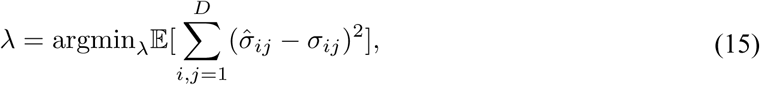

where σ*_ij_* is the *ij*-th entry in the corresponding covariance matrix **Σ**. This equation has an analytic solution. Let **Z** be the variance over training trials of the outer product of the vector of residuals, i.e. if **R** = **Y** - **BW**, then

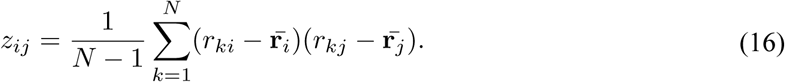

Then the solution for *λ* is

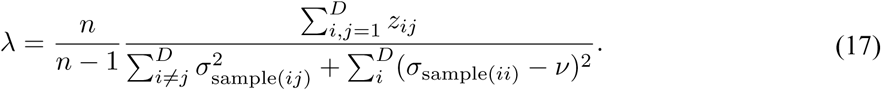

##### 2b Tuning-similarity-weighted (TSW) shrinkage (following TAFKAP)

We also tested a nonstandard approach to covariance shrinkage developed by the TAFKAP authors, which proved relatively successful for uncertainty decoding in fMRI (van Bergen & Jehee, 2021). Instead of shrinking towards a *D×D* diagonal matrix, this approach involves shrinking towards a more complex target **Σ**_0_. Let **τ** = diag(**Σ**_sample_) be the vector containing each channel’s individual variance. Then, for all *i*, the *i*-th diagonal element of **Σ**_0_ is given by

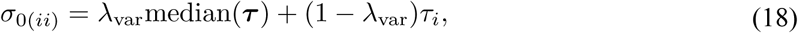

and for all *i* ≠ *j*, the *ij*-th off-diagonal element of **Σ**_0_ is given by

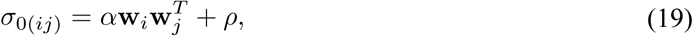

where **w***_i_* is the *i*-th row of the weight matrix **W**. The term **w***_i_***w***_j_^T^* approximates the tuning similarity between the *i* and *j* channels, allowing the covariance structure to take into account tuning-correlated noise (van Bergen & Jehee, 2018). The final covariance estimate is a weighted sum of the sample covariance and target covariance:

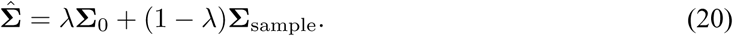

The parameters *α* and *ρ* were fit using ordinary least squares to minimize the squared difference between the off-diagonal elements of **Σ**_0_ and those of the sample covariance:

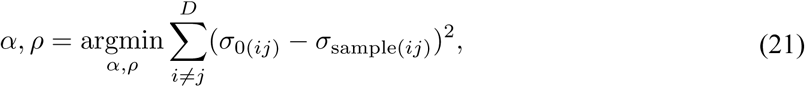

and the hyperparameters *λ* and *λ*_var_ were fit using 4-fold cross-validation within the training data which maximized the average log-likelihood of the resulting covariance estimate given the data in the held-out fold. Both *λ* and *λ*_var_ were fit separately for each time point, for each participant, but outside of the bagging loop.

##### 2c Factor analysis

Since EEG channels can be highly collinear based simply on their spatial proximity, we also tested a factor analysis estimator for noise covariance, which gives a low-rank approximation for the off-diagonal elements of the covariance structure. This approach is based on a generative model where the *D*-dimensional vector of channel residuals **r** = **y** – ***f***(θ) is related to an *L*-dimensional vector of independent latent variables **v**, with *L* < *D*:

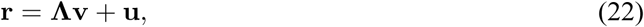

where **Λ** is a *D×L* matrix of loadings and **u** is independent noise, i.e. **u** ∼ N (**0**, **Ψ**), where **Ψ** is *D×D* and diagonal. We also assume that the latent variables are i.i.d. standard normal, i.e. **v** ∼ N (**0**, I*_L_*). It follows that

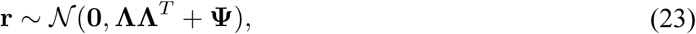

so this model yields a noise covariance estimate **Σ^^^** = **ΛΛ***^T^*+ **Ψ**. To fit **Λ** and **Ψ** we used an expectation maximization approach (Ghahramani & Hinton, 1996). On the *m*+1-th iteration of the algorithm, we updated our parameter estimates with the solutions which maximize the expectation (with respect to the distribution of **v**|**r**, **Λ***_m_* **Ψ***_m_*) of the joint log-likelihood of **r** and **v**.

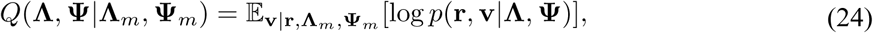

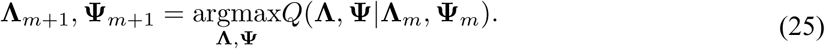

This is solved by

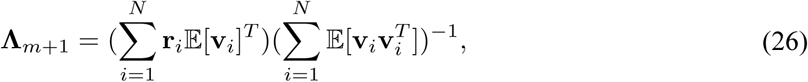

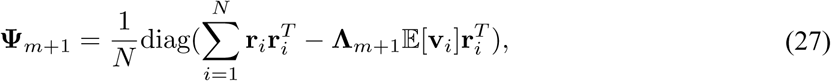

where

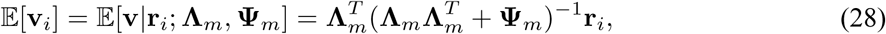

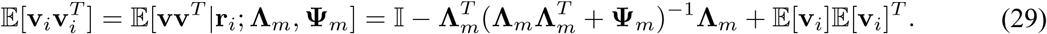

We initialized the expectation maximization algorithm by setting **Λ**_0_ to a *D×L* matrix whose columns are the first *L* eigenvectors of **Σ**_sample_ scaled by the square roots of their respective eigenvalues, and by setting **Ψ**_0_ to I*_D_* ⊙ **Σ**_sample_ - **Λ**_0_**Λ**_0_*^T^*, where ⊙ refers to the elementwise product. We ran the algorithm until the elementwise Euclidean distance between two consecutive covariance estimates was less than 10^-4^.

In order to fit the hyperparameter *L*, we used a 4-fold cross-validation procedure within the training data which maximized the average log-likelihood of the resulting covariance estimate given the data in the held-out fold. *L* was fit separately for each time point, for each participant, but outside of the bagging loop.

##### 2d Independent variance

To test the importance of estimating between-channel covariance for probabilistic decoding in EEG, we used a covariance matrix where all the off-diagonal elements are set to zero, and the *i*-th diagonal element is the sample variance of channel *i*. This is equivalent to assuming that noise is independent between EEG channels. The covariance matrix estimate is thus given by

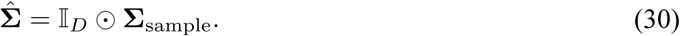

##### 2e Identity matrix

To test the importance of the covariance term for probabilistic decoding in EEG, we used a method where we replaced the covariance matrix with the identity matrix, effectively removing it from the formula for the log-posterior:

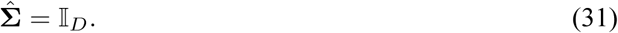

#### Model testing

To characterize model performance, we estimated posteriors over target location for each trial at each time point, using leave-one-out cross-validation so that each trial’s posterior was computed using a model trained on all other trials. We took the circular mean of the posterior to obtain a location estimate from the decoder, and we took the circular variance of the posterior to obtain an uncertainty estimate.

#### Support vector regression

To estimate a benchmark for achievable decoding accuracy on this dataset, we used an established support vector regression (SVR) decoding approach as a baseline (Drucker et al., 1996). For each participant, at each time point, we used the *fitrlinear* function in MATLAB’s Statistics and Machine Learning toolbox, which fits a ridge-regularized SVR model with cross-validation over regularization parameters, to regress both the sine and cosine of the target location onto the preprocessed EEG data. The predicted target location for each test trial is thus given by

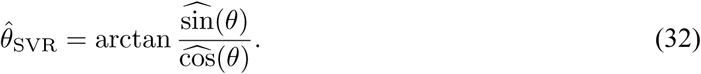

#### Statistical analysis

After computing location estimates and decoded uncertainty for each trial at each time point, we next established the validity of the decoder by comparing the error of location estimates to chance. To do this, we first used a one-sample t-test (2-tailed, *α* = 0.05) to compare the mean absolute error across subjects to the expected mean absolute error of a completely random decoder (in this case, 90°) at each time point. Then, to determine time windows in which decoding was significantly above chance, we used a nonparametric cluster correction approach (Maris & Oostenveld, 2007), in which we identified clusters of consecutive timepoints where the t-statistic crossed the *α* = 0.05 threshold and took the sum of all t-statistics within that cluster. Then, to determine the *α* = 0.05 significance threshold at the cluster level, we generated a null distribution by shuffling the stimulus location labels 500 times and repeating the decoding procedure on the shuffled data, then taking the maximum cluster t-statistic within each permutation.

We benchmarked the precision of the decoded location estimates against existing methods by comparing the mean error of the location estimate from the probabilistic decoder to that from the SVR decoder at each time point using a paired t-test (2-tailed, α = 0.05), followed by permutation-based cluster correction, using null distributions generated by shuffling the decoding method labels, to determine significant time windows.

To test the relation between decoded error and decoded uncertainty, we calculated the trial-by-trial Pearson correlation between the decoder error and decoded uncertainty at each time point and compared the correlation coefficients to zero. Finally, we tested the functional relevance of the decoded uncertainty metric by calculating the trial-by-trial Pearson correlation between the reported arc length and the decoded uncertainty at each time point and comparing the correlation coefficients to zero. For these correlation analyses, we tested for significant differences from zero by Fisher z-transforming the correlation coefficients and performing a z-test (2-tailed, α = 0.05) with permutation-based cluster correction using null distributions generated by shuffling the stimulus location labels 500 times and repeating the decoding procedure on the shuffled data.

#### Covariance estimator comparison

While the probabilistic decoding analyses we performed on the full time series used the standard shrinkage covariance estimator, we also tested decoding performance in a time window around the peak decoding accuracy for four additional covariance estimators. These were the TSW shrinkage and factor analysis estimators, which were selected to better account for tuning-correlated noise and low-rank correlation structures, respectively; and the independent variance and identity matrix estimators, which were used to test how essential the noise covariance estimate is for accurate decoding. To summarize the performance of each model variant, we identified a time window in which peak decoding accuracy was achieved for most participants (150-250 ms) and averaged the decoding error within that period to obtain an estimate of model accuracy. We also averaged the correlation coefficient between decoding error and decoded uncertainty within the same peak time window to determine the trial-by-trial relation between decoded uncertainty and decoding performance. Finally, we averaged the correlation coefficient between decoded uncertainty and reported uncertainty within the peak time window to assess the functional relevance of decoded uncertainty from each model variant. We first compared each of these metrics to chance (90° in the case of absolute error, zero in the case of both correlations) for each model using one-sample t-tests. Then, we compared the metrics across model variants using a one-way repeated-measures ANOVA, following up on significant ANOVA results with post-hoc Bonferroni-corrected paired-sample t-tests.

#### Testing informativeness of single-trial uncertainty estimates versus single-trial decoding error

Although average decoding error across trials is a useful measure of the overall reliability of stimulus information encoded in a set of brain data, it is theoretically unreliable at the single-trial level because it is randomly distributed when information is low. As a result, even when no stimulus information is available, error on a given trial can be arbitrarily low just by chance. We hypothesized that decoded uncertainty would not have this issue, making it more informative than decoding error as a trial-by-trial measure of stimulus information. To assess this hypothesis, we compared the trial-by-trial distribution of each measure in pre-versus post-target time points, which served as a proxy for the absence versus presence of stimulus information.

To quantify the separability of the pre- and post-target distributions for each measure, we used a receiver operator characteristic (ROC) approach. For each participant, we extracted the decoded uncertainty from each trial at a time point 100 ms prior to target presentation, and then we determined the time point where average decoding error was lowest for that participant (mean across participants: 222 ms post-target onset, SD = 29 ms) and extracted the decoded uncertainty from each trial at that time point. We then used the *perfcurve* function in MATLAB to calculate the area under the ROC curve (AUC) for a binary classifier determining whether a decoded uncertainty measurement was from the pre- or post-target distribution. We repeated the same analysis for decoding error instead of decoded uncertainty. We tested for significant differences between AUCs from the decoded uncertainty distributions and those from the decoding error distributions using a bootstrapping procedure. Across 1000 bootstrap iterations, we resampled trials with replacement for each participant, subtracting the average AUC for decoded uncertainty from that for decoding error. Then we computed the 95% confidence interval from the resulting distribution of differences.

## Results

We recorded EEG while participants estimated the angular location of a brief, low-contrast radial grating target presented at a fixed eccentricity and random polar angle on each trial. Participants also reported subjective uncertainty about their location estimate on each trial by adjusting the width of an arc around their estimate. Participants earned points if the true location of the target fell within the arc, and they gained more points for smaller arcs (Li et al., 2021). This points system incentivized participants to choose a smaller arc when they were more certain about their location estimate. We then fit a probabilistic decoding model to the EEG time series data to determine whether information about the stimulus location and subjective uncertainty were represented in the topography of the EEG signal across time. We examined the time courses of decoding error and decoded uncertainty, assessed their trial-by-trial correlation, and tested whether decoded probability distributions predicted behavioral responses.

### Subjective uncertainty reports relate to behavioral error

We first evaluated participants’ behavior in the location estimation task to determine their metacognitive sensitivity. To do so, we tested whether there was a trial-by-trial relation between subjective uncertainty (arc length) and objective performance (response error; Figure 3A). At the group level, arc lengths were wider when the absolute error of the corresponding location estimate was greater (Figure 3B, mean r = 0.393, p < 0.001), demonstrating that participants had metacognitive sensitivity to their perceptual uncertainty about the stimulus location.

**Figure 3.**
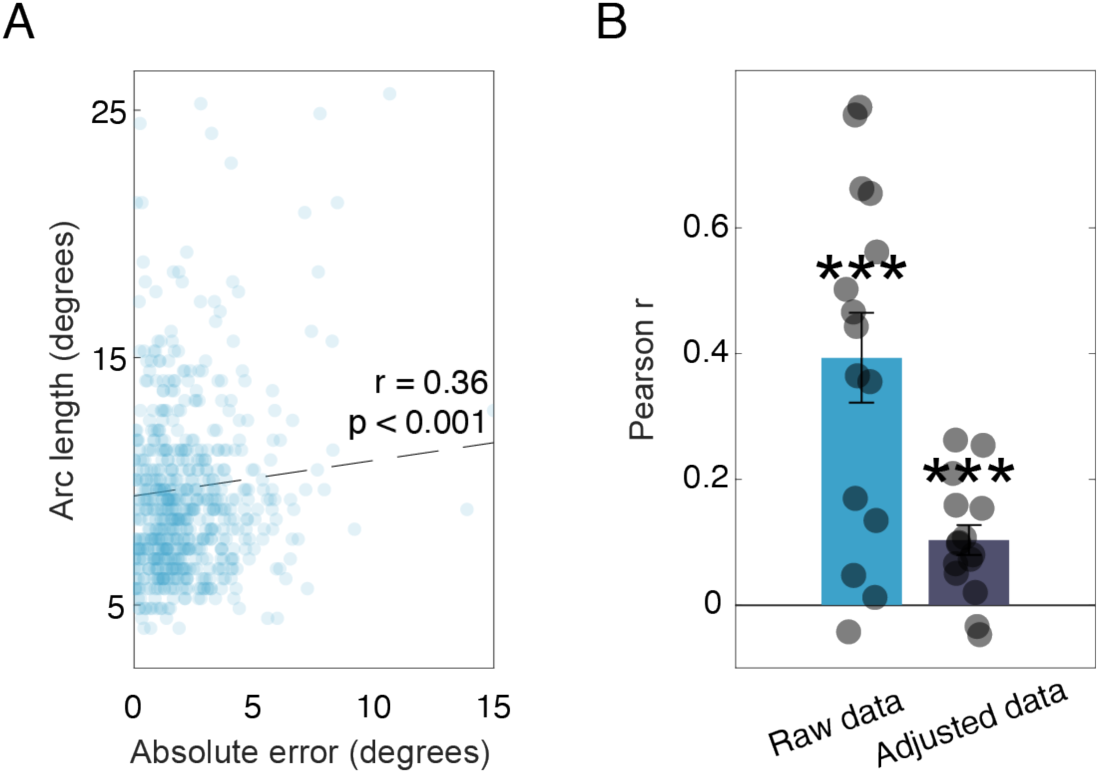
Behavioral uncertainty reports track performance accuracy. A) Relation between arc length and absolute error for one example participant. Each dot is one trial. The dashed line shows the best linear fit. B) Mean correlation coefficient between absolute error and arc length before (raw data) and after (adjusted data) excluding likely miss trials and regressing out linear time-on-task effects. Correlation is significantly above zero in both cases. Each dot is one participant. Bars show means and error bars show +/- 1 sd across participants. *** p < 0.001.

To ensure that this correlation was not driven entirely by “miss trials,” on which location information was completely lost, or time-on-task effects such as learning, we corrected for these factors and repeated the analysis (see Methods). A significant positive correlation between arc length and absolute error remained even after excluding the trials that were likely to be misses and regressing out time-on-task effects (Figure 3B, mean r = 0.1035, p < 0.001). Metacognitive sensitivity in this task is therefore not driven only by a few difficult trials or by practice effects.

### Decoded uncertainty tracks decoding error across trials and time

After establishing metacognitive sensitivity behaviorally, we examined the output of the probabilistic decoder. First, we evaluated the decoder’s performance accuracy. We used the mean of the decoded probability distribution at each time point and each trial as the decoder’s prediction of stimulus location, and we quantified trial-by-trial decoding error for each time point by computing the absolute circular distance between the prediction and the true target location. We evaluated several different variants of the probabilistic decoder; the main time series results we report all used the standard shrinkage decoder, which we found balanced accuracy and computational efficiency.

We first examined the time courses of decoding error and decoded uncertainty. Decoding performance remained around chance during the pre-stimulus period before increasing, as indicated by a reduction in error, shortly after stimulus onset (Figure 4A). The probabilistic decoder performed significantly above chance starting at 160 ms (p < 0.05, cluster corrected), and it continued to perform above chance through the end of the epoch at 500 ms. To test whether the probabilistic approach sacrificed sensitivity relative to standard decoding approaches, we also compared decoding error to a benchmark support vector regression (SVR) method. The performance of the SVR decoder and the probabilistic decoder did not differ significantly throughout the epoch after cluster correction, demonstrating that probabilistic decoding in EEG performs indistinguishably from standard decoding methods (Figure 4A).

**Figure 4.**
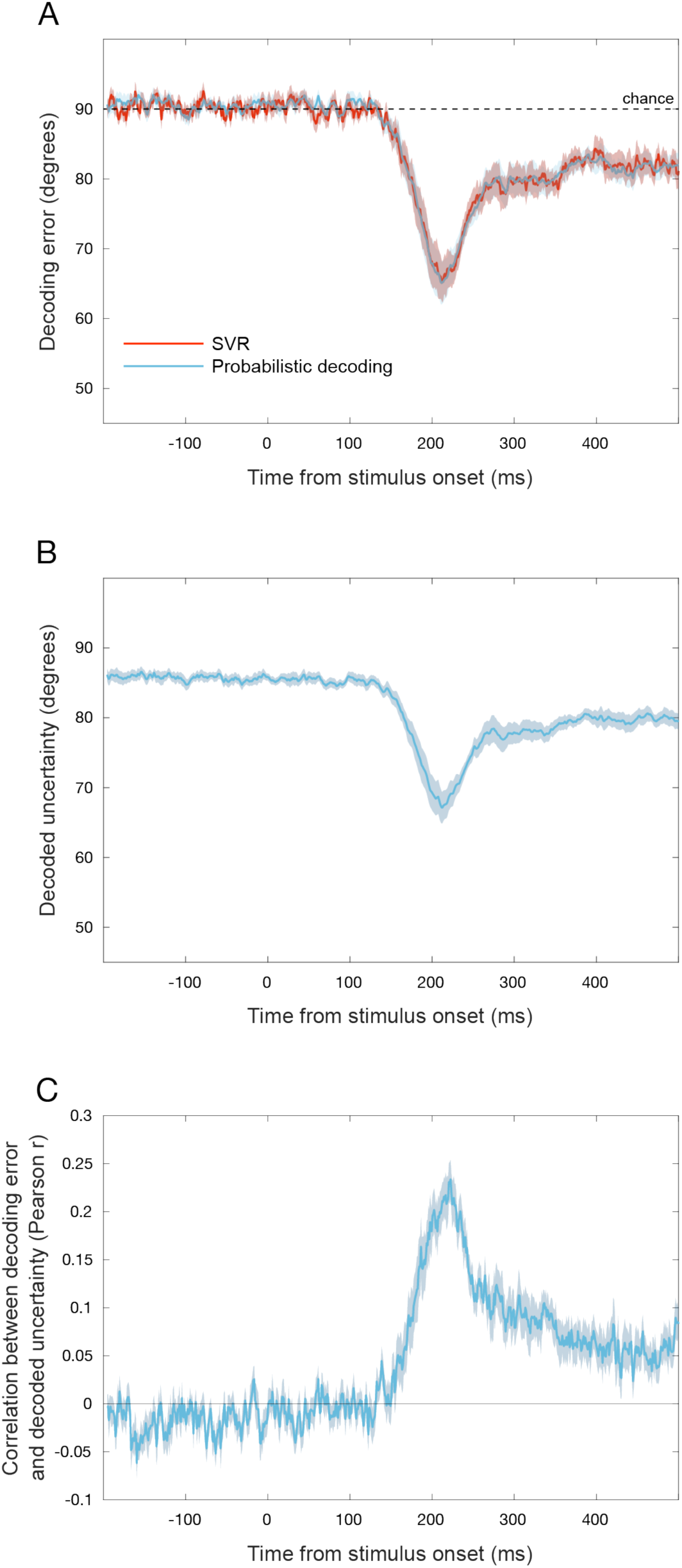
Decoding time series. A) Absolute error from the probabilistic decoder and a benchmark support vector regression decoder across time. No significant differences between SVR and probabilistic decoding. Chance line at 90° shows expected absolute error of a decoder with random outputs. B) Decoded uncertainty across time. C) Trial-by-trial correlation between decoded uncertainty and decoding error across time. Time series show group means (n = 15) with +/-1 SEM error ribbons.

In addition to performing as well as standard decoding methods in terms of accuracy, probabilistic decoding has the advantage of providing a measure of decoding uncertainty. Here, we leveraged EEG’s high temporal resolution to characterize the dynamics of this decoded uncertainty, using the circular variance of the decoded probability distribution at each time point and each trial as a time-resolved measure of decoded uncertainty. We first explored the time course of decoded uncertainty averaged across trials (Figure 4B). Decoded uncertainty had a remarkably similar time course to decoding error: across time, each participant’s trial-average decoding error and trial-average decoded uncertainty were highly correlated (mean Pearson r across participants: 0.965, SD = 0.030). This correlation suggests that decoded uncertainty decreases when location information becomes available in the EEG signal, as expected.

We then tested whether decoded uncertainty tracked the reliability of stimulus information in the EEG signal at a trial-by-trial level. To do so, we assessed whether decoded uncertainty at a given time point had a significant positive correlation with decoding error at the same time point, across trials (Figure 4C). We found that this correlation was significant starting at 178 ms (p < 0.05, cluster corrected), indicating that decoded uncertainty is lower on trials where more location information is available in the EEG signal. While trial-average decoding error and decoded uncertainty were closely related across time, the peak trial-by-trial correlation between these measures reflects a relatively modest relationship (r = 0.256), indicating that the two measures may have picked up non-overlapping information from the underlying EEG signal.

### No evidence for a relation between decoder estimates and behavior

We next asked whether the probabilistic decoder could predict behavioral responses. For each time point, we tested whether decoding error correlated trial-by-trial with behavioral response error (Figure 5A,B), as well as whether decoded uncertainty correlated trial-by-trial with reported uncertainty (arc length) (Figure 5C,D). However, we found no significant clusters for either analysis.

**Figure 5.**
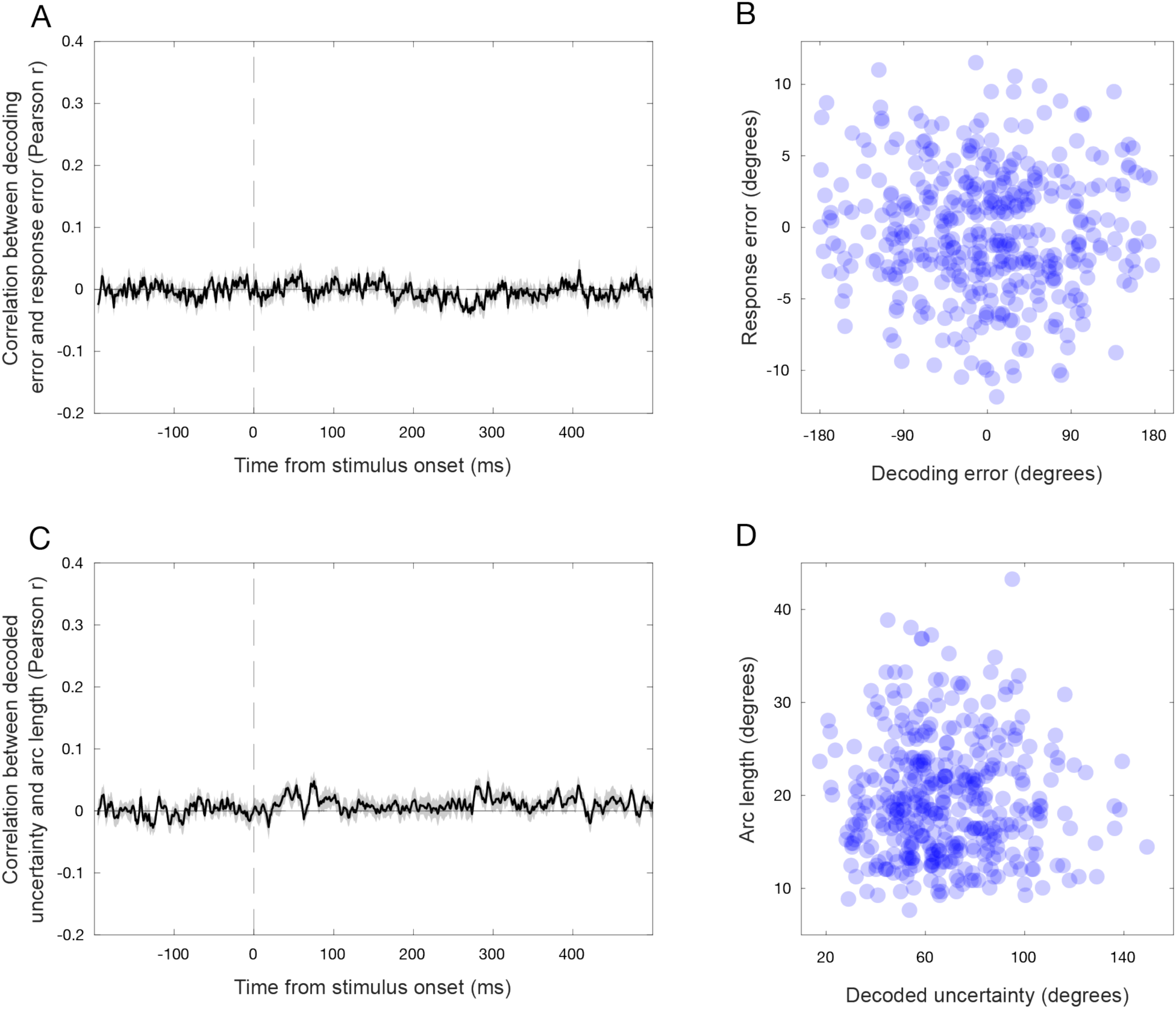
Brain-behavior correlation time series. A) Trial-by-trial correlation between signed decoding error and signed response error. B) Example scatterplot showing the relationship between response error and decoding error at 200 ms for one participant. Each dot is one trial. C) Trial-by-trial correlation between decoded uncertainty and arc length responses. D) Example scatterplot showing the relationship between arc length and decoded uncertainty at 200 ms for one participant. Each dot is one trial. Neither correlation is significantly different from zero at any time point after cluster correction. Time series show group means (n = 15) with +/-1 SEM error ribbons.

### The noise covariance structure impacts probabilistic decoding performance but not behavioral correlations

While the core principles behind probabilistic decoding are theoretically applicable to any neural recording method, estimating the noise covariance structure may require different approaches for different types of data. The noise covariance model effectively allows the decoder to discount patterns in the data that are likely to arise from noise rather than stimulus information, and the TAFKAP authors noted that using a different noise covariance estimator could produce dramatic differences in decoder output from fMRI data (van Bergen & Jehee, 2018). Because EEG data has different noise properties than fMRI data, the impact of the noise covariance model on probabilistic decoding for EEG must be determined empirically.

We tested a variety of methods for estimating the noise covariance of EEG data and compared the decoding results across methods. Three methods accounted for correlated noise across EEG channels: the standard shrinkage method, which is an established approach for EEG data (Blankertz et al., 2011) and was used for the time series analyses above; the tuning-similarity-weighted (TSW) shrinkage method used in TAFKAP (van Bergen & Jehee, 2021), which is designed to account for tuning-correlated noise; and the factor analysis method, which gives a low-rank approximation for the between-channel covariance structure. We also tested two additional methods to evaluate the overall importance of the noise covariance estimate in probabilistic decoding: an independent variance model and an identity matrix model, neither of which allow the decoder to account for correlated noise. To summarize model performance, we averaged decoding error for each model within a key time window (150-250 ms post-target onset, Figure 6A).

**Figure 6.**
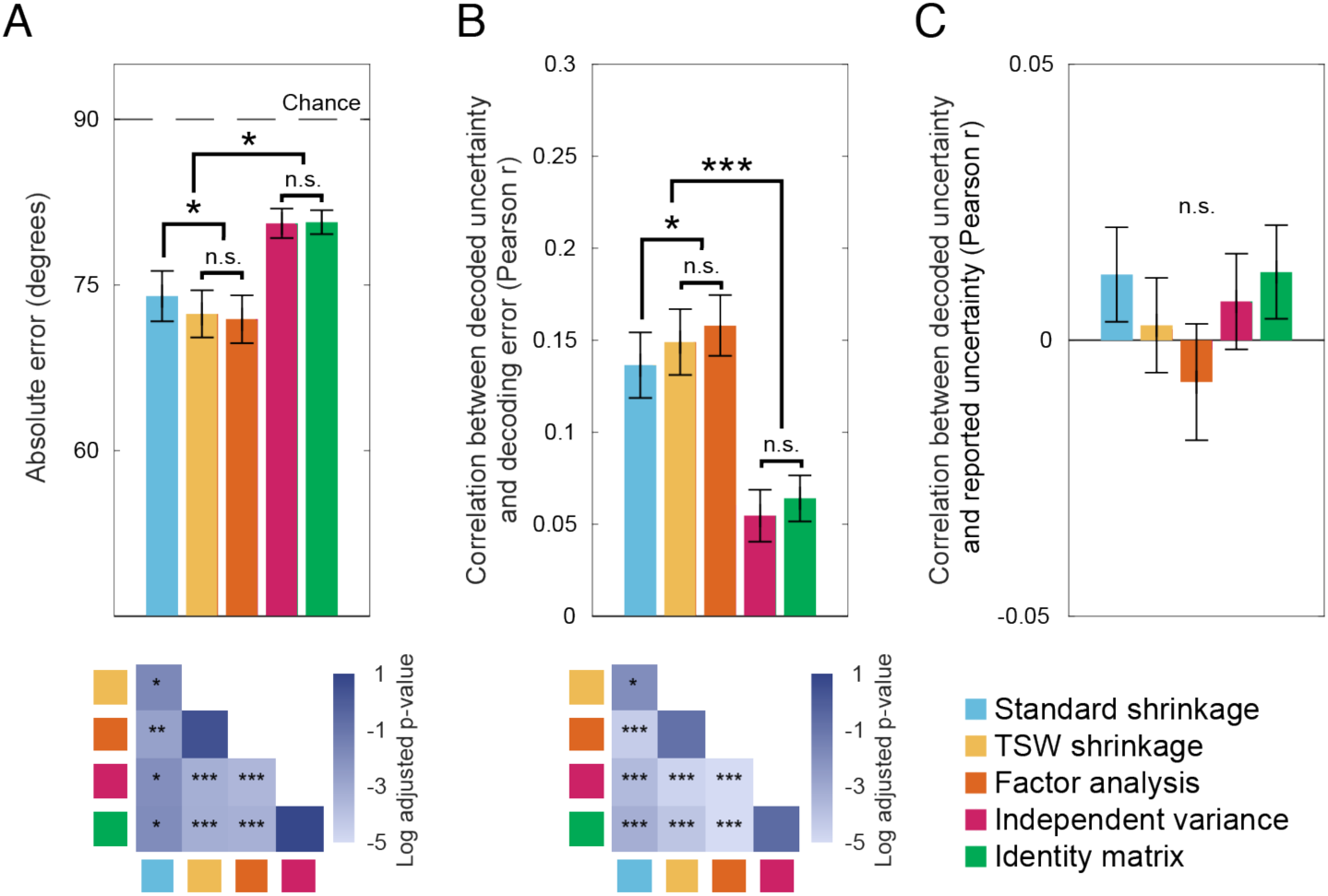
Comparison among different noise covariance estimators. Bars show mean across participants and error bars show +/- 1 SEM. All values are averaged from timepoints 150-250 ms after stimulus onset. A) Absolute decoding error for each covariance method. Error is significantly below chance for all methods. Lower plots show p-values for pairwise comparisons after Bonferroni correction. Stars above brackets reflect the highest p-value among relevant pairwise comparisons. B) Trial-by-trial correlation between decoded uncertainty and decoding error for each covariance method. Correlation is significantly above chance for all methods. Same conventions as in panel A. C) Trial-by-trial correlation between decoded uncertainty and reported arc length for each covariance method. Correlation is not significantly different from chance for any method. * p < 0.05, ** p < 0.01, *** p < 0.001.

All models showed location decoding, but including between-channel correlations in the noise covariance improved performance. For all five methods, decoding error was significantly below chance-level (all t(14) > 7.007, all p < 0.001), showing that regardless of the covariance estimate, probabilistic decoding of EEG data provides above-chance predictions. However, average decoding error differed significantly between the five covariance methods (repeated-measures ANOVA, F(4, 56) = 25.958, p < 0.001). The three covariance methods that allowed for between-channel correlations (standard shrinkage, TSW shrinkage, and factor analysis) performed significantly better than the two that did not allow for any between-channel correlations (independent variance and identity matrix) (paired-samples t-tests Bonferroni-corrected for 10 comparisons, all t(14) > 3.966, all adjusted p < 0.014), showing that the off-diagonal elements of the noise covariance term were important for more accurate decoding. There was no significant difference between the decoding error for the independent variance and identity matrix methods (t(14) = 0.289, adjusted p = 1.00).

Among models with between-channel correlations, the specific noise covariance structure mattered for performance. Standard shrinkage was slightly but significantly outperformed by both TSW shrinkage (t(14) = 3.798, adjusted p = 0.02) and factor analysis (t(14) = 4.963, adjusted p < 0.01), which were not significantly different from each other (t(14) = 1.074, adjusted p = 1.00), showing that more sophisticated noise covariance models can lead to more accurate decoding. However, we note that, due to the cross-validation involved in fitting the covariance models, TSW shrinkage and factor analysis increased overall computing time by approximately 50 and 10 times, respectively, relative to standard shrinkage.

The choice of noise covariance also impacted the relation between decoded uncertainty and decoding error. To assess the reliability of the decoded uncertainty measure derived from each covariance method, we averaged the trial-by-trial correlation between decoding error and decoded uncertainty within the same key window (Figure 6B). For all five methods, this correlation was significantly above chance (all t(14) > 3.877, all p < 0.002), showing that decoded uncertainty can be meaningful even with a simplified model. Still, the correlation differed significantly between the five covariance methods (F(4, 56) = 52.191, p < 0.001). Similar to decoding error, the correlation was stronger for standard shrinkage, TSW shrinkage and factor analysis than it was for independent variance or the identity matrix (all t(14) > 5.635, all adjusted p < 0.001), while independent variance and the identity matrix did not significantly differ from each other (t(14) = 2.431, adjusted p = 0.291), indicating that estimating between-channel correlation is also important for generating meaningful decoded uncertainty metrics. Additionally, the correlation for standard shrinkage was slightly but significantly weaker than that for both TSW shrinkage (t(14) = 4.012, adjusted p = 0.012) and factor analysis (t(14) = 7.319, adjusted p < 0.001), while TSW shrinkage and factor analysis were not significantly different from each other (t(14) = 2.743, adjusted p = 0.159), further demonstrating the benefits of these more sophisticated covariance estimators.

Finally, we assessed the functional relevance of decoded uncertainty measures derived from each noise covariance model by averaging the trial-by-trial correlation between decoded uncertainty and reported arc angle within the same key time window (Figure 6C). This correlation was not significantly different from zero for any of the covariance methods (all t(14) < 1.458, all p > 0.168). There were also no significant differences in this correlation between any of the methods (F(4,56) = 1.466, p = 0.224). Thus, despite the methods showing differences in their decoding performance and the reliability of decoded uncertainty as a measure of information in the brain signal, none of the methods yielded decoded uncertainty measures that were related to subjective uncertainty.

### Decoded uncertainty is more informative than decoding error on a single trial

Finally, to characterize the reliability of both decoded uncertainty and decoding error as trial-by-trial measures of stimulus information, we asked whether these measures consistently differentiated between time points where a stimulus had or had not been presented at a single-trial level. To do so, we assessed how well the trial-by-trial distribution of error and uncertainty could be separated into pre-versus post-target time points (Figure 7). For error, the distribution across trials had a folded bell shape for the post-target time point (Figure 7A), where errors were more likely to be close to zero due to the decoder’s above-chance performance. In contrast, the error distribution was approximately uniform for the pre-target time point, reflecting the fact that when there is no stimulus information, the decoder’s prediction is random with respect to the true target location. For uncertainty on the other hand, the distribution was bell-shaped for both the pre- and post-target time points, and the mean of the pre-target distribution was higher than that of the post-target distribution (Figure 7B). This pattern shows that when there is no stimulus information, the decoded uncertainty is reliably high.

**Figure 7.**
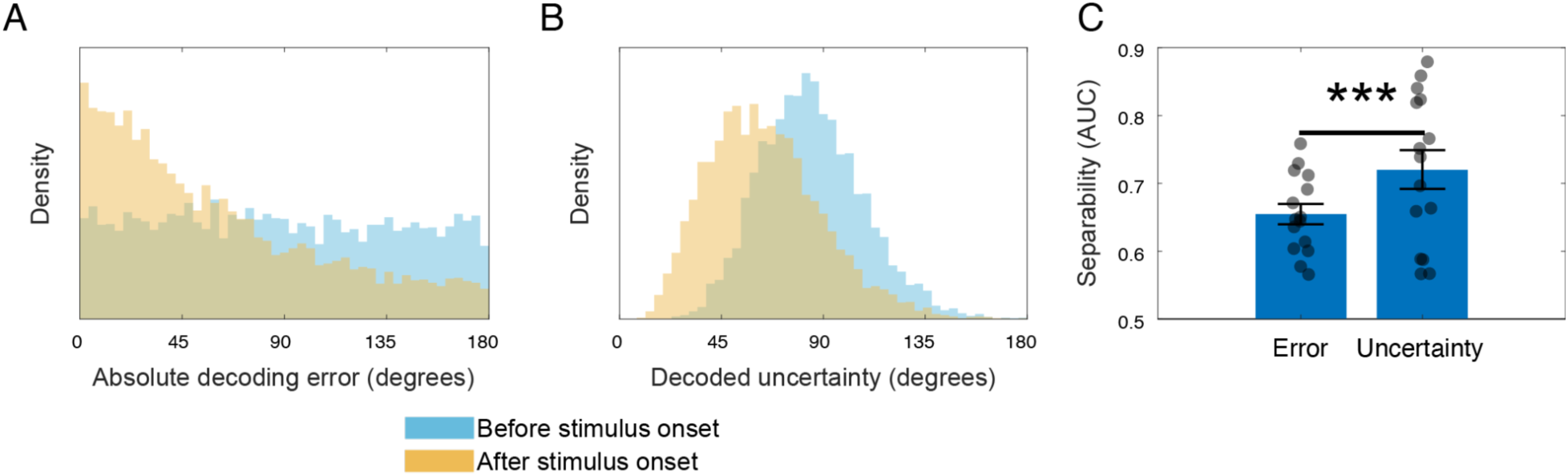
Decoded uncertainty is superior to decoding error as a trial-by-trial indicator of stimulus information. Histograms show distributions combined across all participants, whereas ROC analysis reflects within-subject classification of pre-versus post-target time points. A) The distribution of decoding error 100 ms prior to stimulus onset versus at the time point after stimulus onset where decoding accuracy is the highest for that participant. B) The distribution of decoded uncertainty 100 ms prior to stimulus onset versus at the time point after stimulus onset where decoding accuracy is the highest. C) The area under the ROC curves for the classification of pre-versus post-target measurements using decoding error or using decoded uncertainty. Higher AUC for decoded uncertainty means that the two uncertainty distributions are more separable than the two error distributions. Each dot is one participant. Bars show means and error bars show +/- 1 SEM across participants (n = 15). *** p < 0.001.

We quantified the separability of the pre-target and post-target distributions for each decoding measure by considering two possible binary classifiers: one that predicts whether a decoding error value was taken from a pre- or post-target time point and another that predicts whether a decoded uncertainty value was taken from a pre- or post-target time point. For each participant, we computed the ROC curves for these two classifiers and took the AUC (Figure 7C). Across participants, AUCs were significantly higher for the decoded uncertainty classifier (mean difference: 0.066, bootstrap 95% CI: [0.0515, 0.0799], p<0.001). Thus decoded uncertainty was superior to decoding error as a trial-by-trial indicator of the presence of stimulus information.

## Discussion

We developed an approach for the time-resolved measurement of stimulus uncertainty in the human brain to address the limited temporal resolution of existing code-driven metrics developed for fMRI. Our approach, adapted from TAFKAP (van Bergen & Jehee, 2021), fits generative models for neural data at each time point, allowing a time-varying probability distribution over a target feature to be decoded on each trial.

We developed and tested the method in the context of stimulus location processing in EEG. The results showed that time-resolved uncertainty about stimulus location could be decoded from EEG and reflected the stimulus information available on a single trial, though with no apparent relation to behavioral uncertainty reports. An accompanying toolbox (PDAT: Probabilistic Decoding Across Time) allows researchers to investigate the dynamics of uncertainty inherent to time-varying brain signals over the course of stimulus processing.

Compared to TAFKAP for fMRI, we used the same tuning model and general approach, but we tested several adjustments to the noise covariance model. Modelling the noise covariance using a method that is standard for EEG produced large improvements in accuracy compared to models that did not consider noise covariance. Accuracy showed further slight improvements with models that make assumptions about the underlying structure of noise correlations, including TAFKAP, though at a cost to computational efficiency.

We first demonstrated the feasibility of probabilistic decoding for EEG, showing that decoding full probability distributions over stimulus location trial-by-trial produced comparable accuracy to a standard point-estimate decoder while also providing a time-varying measure of decoded uncertainty about the stimulus location. Importantly, decoded uncertainty correlated with decoding error across trials at each time point, as long as decoding performance was above chance. This correlation across trials occurred even though all stimuli were physically identical except for their location. These findings show that decoded uncertainty is a meaningful trial-by-trial measure of the stimulus information available in the EEG signal at each time point.

We also tested the decoded uncertainty measure to determine to what extent it reflects a neural representation of stimulus uncertainty. Such tests are important because decoded uncertainty may reflect a combination of neural noise and other noise sources—like noise in the physiological measurement—not related to stimulus processing. The interpretation of decoded uncertainty therefore depends on which sources of noise contribute to it.

It has been proposed (Pohl et al., 2026; Walker et al., 2023) that evidence for a given signal being a neural representation of uncertainty can be categorized under four criteria. 1) Sensitivity: changes in the representation of uncertainty should be related to changes in sensory uncertainty. 2) Specificity: the relationship between sensory uncertainty and the uncertainty representation should remain even when controlling for possible confounds, such as stimulus manipulations. 3) Invariance: the relationship between sensory uncertainty and the uncertainty representation should occur even across levels of other variables. 4) Functionality: the uncertainty representation should influence behavioral responses. Typically, a single experiment does not address all criteria (Walker et al., 2023). Here, we focused on sensitivity, specificity, and functionality, and did not address invariance.

One approach to test for sensitivity is to vary a stimulus parameter, like contrast, that is expected to affect uncertainty, and assess how a candidate measure of uncertainty varies with that stimulus manipulation (Orbán et al., 2016; Walker et al., 2020). In this experiment, however, we held constant stimulus variables that might affect uncertainty to test whether we could decode uncertainty arising solely from neural variability, rather than from physical stimulus properties, allowing strong tests of specificity and functionality (Geurts et al., 2022; Li et al., 2021).

To assess sensitivity, then, we used the trial sequence as a proxy for sensory uncertainty, as we can be sure that the brain signal contained no stimulus information before the target appeared. We found that decoded uncertainty followed the time course of decoding error, showing that the uncertainty metric was sensitive to the timing and degree of stimulus information. Moreover, decoded uncertainty was a more informative indicator of pre-vs. post-target time points than was decoding error. Thus, while one could use decoding error to approximate uncertainty, decoded uncertainty is a more sensitive uncertainty metric, which constitutes an advantage of probabilistic decoding over standard point-estimate decoding approaches.

To assess specificity, we showed that decoded uncertainty correlates on a trial-by-trial level with decoding error, even as the stimulus remained constant across trials. Note that such a relationship could arise due to common sensory noise, common measurement noise, or some combination, so whether or to what degree decoded uncertainty reflects neural uncertainty cannot be determined from trial-by-trial correlations alone. Further work exploring how the decoded uncertainty obtained from this method is affected by various stimulus manipulations would provide additional evidence regarding its sensitivity and specificity.

Finally, we tested but found no evidence for the functionality of the decoded uncertainty in EEG, as it had no significant trial-by-trial correlation with reported uncertainty in the behavioral location estimation task. This finding diverged from those of experiments that used TAFKAP with fMRI, which did find a correlation between decoded uncertainty and reported uncertainty. Such correlations have been observed for both orientation reports (Geurts et al., 2022) and location reports using a similar task to the current study but with a long working-memory delay (Li et al., 2021). The current results thus suggest that probabilistic decoding with EEG may offer limited insight into metacognition, though it will be important in future work to test for functionality using other stimuli and tasks.

Several differences between fMRI and EEG may contribute to the lack of brain-behavior correlations for TAFKAP adapted for EEG, despite positive findings in fMRI. The lower signal-to-noise ratio in EEG may contribute to this null effect, as well as the better spatial resolution of fMRI and the ability to select voxels that respond to the stimulus. The TAFKAP model may also better capture stimulus dependencies in fMRI than EEG data, as indicated by the findings that TAFKAP yields more accurate decoding than SVR for fMRI (van Bergen & Jehee, 2021) but not for EEG.

Differences in the temporal properties of EEG and fMRI may also contribute to brain-behavior correlation differences. First, as the temporal resolution of the fMRI BOLD response is on the order of seconds, the brain-behavior correlations observed in fMRI could arise from later signals not captured in our 500 ms epoch. Contributing to this possibility, the fMRI studies had delays of at least 6 seconds after the stimulus presentation until participants could make their behavioral reports (Geurts et al., 2022; Li et al., 2021). Indeed, Geurts et al. (2022) reported that the relationship between decoded uncertainty and reported confidence emerged later in the working memory retention interval than did above-chance decoding. It is therefore possible that the functional element of neural uncertainty is represented relatively late in stimulus processing or only during the working memory retention interval, which our protocol would not have captured.

Second, the brain-behavior correlation observed in fMRI could arise due to integration of signals across time that is not captured by the fine timescale of EEG. For instance, in sampling-based coding schemes, uncertainty is represented by variability across time within single neurons (Haefner et al., 2016; Orbán et al., 2016; Zhu et al., 2024), which predicts that an instantaneous measure of stimulus information would not contain a functional representation of uncertainty. This principle would favor slower, more integrative neural measures of uncertainty as opposed to those with high temporal resolution.

Despite the lack of evidence for a functional role of time-resolved uncertainty decoded from EEG, the measure is both sensitive and specific to stimulus uncertainty in the EEG signal and provides a single-trial metric of the stimulus information available in the signal. Thus, uncertainty decoding could be a valuable tool as a measure of encoded information across time.

Although decoding error, which can also be derived from standard decoding methods, has frequently been used as a proxy for encoded information (Bae & Luck, 2018; Grootswagers et al., 2017), decoded uncertainty offers advantages over decoding error. As we demonstrated here, decoded uncertainty is a more reliable indicator of stimulus information than decoding error at the single-trial level. This finding can be explained by the fact that decoding error approaches a uniform distribution (i.e., becomes unreliable) as stimulus information is reduced, whereas decoded uncertainty provides similar reliability across the full range of stimulus information levels.

Additionally, a limitation of decoding error is that it can only be calculated in comparison to a known ground truth, whereas decoded uncertainty can be calculated even when the experimenter does not have access to the ground truth for a given test trial, so long as they have the ground truth for trials in the training data. For example, many experiments that use near-threshold stimuli include trials on which no stimulus is presented. On these trials, decoding error does not exist, but decoded uncertainty can be measured, providing information about the amount of random stimulus-like signal in the brain data from that trial. Such an approach could shed light on the neural basis of false alarms and illusory percepts.

## Declaration of the use of AI

We did not use AI in writing this paper.

## Data and code availability

All data and code used in the production of this manuscript are available at https://doi.org/10.5281/zenodo.22099566. Our main decoding analysis MATLAB code is available in a user-friendly format through our open-source Probabilistic Decoding Across Time (PDAT) toolbox: https://github.com/denisonlab/pdat.

## Author contributions

Jeffrey Nestor: Conceptualization, data collection, formal analysis, software toolbox, visualization, writing – original draft, writing – review and editing

Karen J. Tian: Conceptualization, data collection, visualization, writing – review and editing

Angus F. Chapman: Conceptualization, data collection, writing – review and editing

Rachel N. Denison: Conceptualization, writing – review and editing

## Acknowledgements

This research was supported by Boston University start-up funds to R.N.D. as well as by the National Defense Science and Engineering Graduate Fellowship to K.J.T.

